# VDJ-Insights: simplifying the annotation of genomic IG and TCR regions

**DOI:** 10.1101/2025.08.15.670470

**Authors:** Susan E. Ott, Giang N. Le, Sayed J. Mohammadi, Jesse Mittertreiner, Erica M. Pasini, Ronald E. Bontrop, Natasja G. de Groot, Jesse Bruijnesteijn

**Affiliations:** Biomedical Primate Research Centre, Lange Kleiweg 161, 2288 GJ Rijswijk, The Netherlands

**Keywords:** T cell receptor, Immunoglobulin receptor, B cell receptor, annotation tool, V(D)J recombination, CDR1, CDR2, RSS, Human pangenome, annotation software

## Abstract

Accurate annotation of germline immunoglobulin (IG) and T cell receptor (TCR) loci is critical for understanding adaptive immunity. VDJ-Insights provides a user-friendly software package for characterizing these complex immune regions. In addition, it assesses gene segment functionality, identifies recombination signal sequences (RSS), and annotates complementary-determining regions 1 and 2 (CDR1, CDR2). VDJ-Insights achieved over 99% concordance with curated annotations from multiple species, outperforming existing annotation tools. When applied to 95 haplotypes from the Human Pangenome Reference Consortium, VDJ-Insights identified 652 and 275 novel IG and TCR alleles, respectively, highlighting its scalability for large immunogenetic studies.

## Introduction

B cells and T cells are essential components of the adaptive immune system, with their respective receptors, the Immunoglobulin receptor (IG) and the T cell receptor (TCR), serving as direct or indirect mediators of pathogen recognition. Genetically, these receptors are constructed in a similar fashion, involving the somatic rearrangement of variable (V), diversity (D), and joining (J) gene segments during B and T cell development. In B cells, this rearrangement takes place within the heavy (IGH) and light (IGK and IGL) chain regions, while in T cells it occurs across the TRA, TRB, TRG, and TRD regions (1, 2). These genomic regions exhibit substantial diversity, characterized by large structural variations, including deletions, insertions, and duplications of functional segments, as well as by allelic polymorphism (3–8). The overall diversity of human IG and TCR gene segments is well-documented, with 1069 and 1565 human V, D, and J alleles catalogued in databases such as International IMmunoGeneTics Information System (IMGT) and VDJbase (both assessed on 2025-04-30), respectively (9, 10). However, the individual haplotypes of these loci remain largely unexplored due to their complex organization, which hampers assembly, phasing, and annotation using conventional high-throughput methods and thus prevents routine documentation (11, 12).

Understanding and documenting the diversity in these highly complex immune regions is of importance to elucidate the impact of germline variation on antibody and T cell responses. For example, specific IG gene segments have been linked to differential responses to vaccines, including those for malaria and seasonal influenza (13–16). Furthermore, polymorphisms in IGH genes affect the function of neutralizing antibodies in response to HIV and SARS-CoV-2 infection (17–19). In the context of disease and vaccine associations, the IG and TCR loci in biomedically important model species, such as mice and macaques, have also been characterized, though only to a limited extent (11, 20–25). Despite studies across multiple species and the extensive cataloging of human germline alleles in various databases, new sequences continue to be discovered (4, 5, 7, 26, 27). This indicates that even greater diversity is likely to emerge as more individuals and species are examined for their IG and TCR loci. Moreover, while most research to date has focused on coding regions, the non-coding sequences, including the recombination signal sequences (RSS), remain largely unexplored. These non-coding features may be essential for the rearrangement processes that ultimately shape IG and TCR repertoires (28).

With the introduction of long-read sequencing technologies, such as Pacific Biosciences (PacBio) and Oxford Nanopore Technologies (ONT) platforms, the characterisation of novel genomic IG and TCR regions has become more accessible (4, 5, 7). Yet, their assembly and annotation remain time-consuming and require expert curation. To address these challenges, several bioinformatic tools, including gAIRR Suite, IGDetective, and Digger, have been developed to analyse either individual V, D, and J gene segments or the complete genomic IG and TCR regions (29–31). Among these, gAIRR Suite is specifically designed for annotating short-read assemblies, whereas IGDetective identifies gene segments without relying on a predefined segment library (30, 31). The most recently published tool, Digger, enables identification of both known and novel gene segments while also predicting their functionality (29). Building on these advancements, we developed VDJ-Insights, a tool that integrates and enhances similar features by automating the identification and annotation of both coding segments and non-coding elements. Additionally, VDJ-Insights dynamically integrates the latest IMGT-database updates and utilizes IMGT standards to determine functionality of known and newly identified gene segments, assess their Recombination Signal Sequences (RSS), and annotate the Complementary Determining Regions (CDR) 1 and 2 positions (32). The tool supports multi-sample processing and provides a user-friendly web-based interface for data interpretation, visualization, and comparative analysis.

We validated VDJ-Insights by analysing IMGT-curated human sequences, as well as curated sequences from other species such as gorillas, rhesus macaques, and mice (22, 27, 33, 34). Following successful validation, samples from the Human Pangenome Reference Consortium (HPRC) were analysed (35). This analysis resulted in the identification of 652 novel IG and 275 novel TCR V, D, and J alleles, of which 222 IG and 137 TCR were confirmed in at least two individuals. Collectively, these findings highlight the strength of VDJ-Insights in processing large-scale datasets and advancing the identification of both known and novel IG and TCR segments.

## Results

### Overview of the VDJ-Insights workflow

VDJ-Insights is developed to extract, annotate, and analyse the IG and TCR regions from genomic sequences (Fig. 1). First, VDJ-Insights extracts the IG or TCR regions using predefined flanking genes, which serve as markers for identifying the boundaries of the regions of interest. These flanking genes are relatively conserved among species (36, 37). For instance, *RSPH141* and *TOP3B1* flank the *IGL* region in both humans and macaques. Our tool provides default flanking genes for humans and commonly used model species (e.g., rhesus macaque (*Macaca mulatta*) and house mouse (*Mus musculus*); Suppl. Table S1) but also allowing users to define custom flanking genes whenever needed. In cases where an immune region appears fragmented, thus with flanking genes located on distinct contigs, users can initiate an additional scaffolding step within the pipeline to potentially construct complete IG and/or TCR regions (38). Next, a sequence library containing previously reported V, D and J gene segments is mapped to the extracted regions, using Bowtie, Bowtie2, and minimap2 (39–41). When no library is provided by the user, VDJ-Insights automatically retrieves a gene segment library for the specified species from the IMGT database, if available (10). Subsequently, mapped gene segments are filtered and evaluated using BLASTN (42). To determine gene functionality, VDJ-Insights examines multiple criteria, including the assessment of leader sequences, when available, the presence of start codons, the screening for in-frame stop codons, and the evaluation of Recombination Signal Sequences (RSS) (Table 1). In addition, the tool identifies the Complementary-Determining Regions 1 and 2 (CDR1 and CDR2), which represent the hypervariable loops that, together with CDR3, form the antigen binding site of antibodies and TCRs. The germline-encoded CDR1 and 2 are assessed by alignment to their IMGT reference sequences when available. As CDR3 is shaped through somatic recombination, it is currently not included in the VDJ-Insights analysis.

**Table 1.** Functionality criteria for V, D, and J gene segments.

| Gene segment | Functionality criteria | If criterion is not met |
| --- | --- | --- |
| V | Translated protein present | Pseudogene |
|  | Reading frame is a multiple of 3 base pairs | Pseudogene |
|  | Start codon present | Pseudogene |
|  | No in-frame stop codons | Pseudogene |
|  | Stop codon at the end the end of the sequence | ORF |
|  | Donor splice site (GT) present | ORF |
|  | Acceptor splice site (AG) present | ORF |
|  | Minimum two cysteines present | ORF |
|  | Functional RSS motif present | Pseudogene |
| D | Functional RSS motifs present | Pseudogene |
| J | Motifs (Regular Expression: [FW]G.G) present | ORF |
|  | No premature stop codon present | Pseudogene |
|  | Donor splice site (GT) present | Pseudogene |
|  | Functional RSS motif present | Pseudogene |

*Gene segment functionality is classified into three categories: functional, open reading frame (ORF), and pseudogene. Gene segments designated as functional meet all criteria for expression and V(D)J rearrangement. Those classified as ORFs exhibit deviations in rearrangement or splicing signals but maintain an intact open reading frame. Pseudogenes are defined by the presence of frameshift mutations or premature stop codons that disrupt the open reading frame. These functionality classifications are predicted based on genomic features and are not supported by experimental data.*

**Figure 1.**
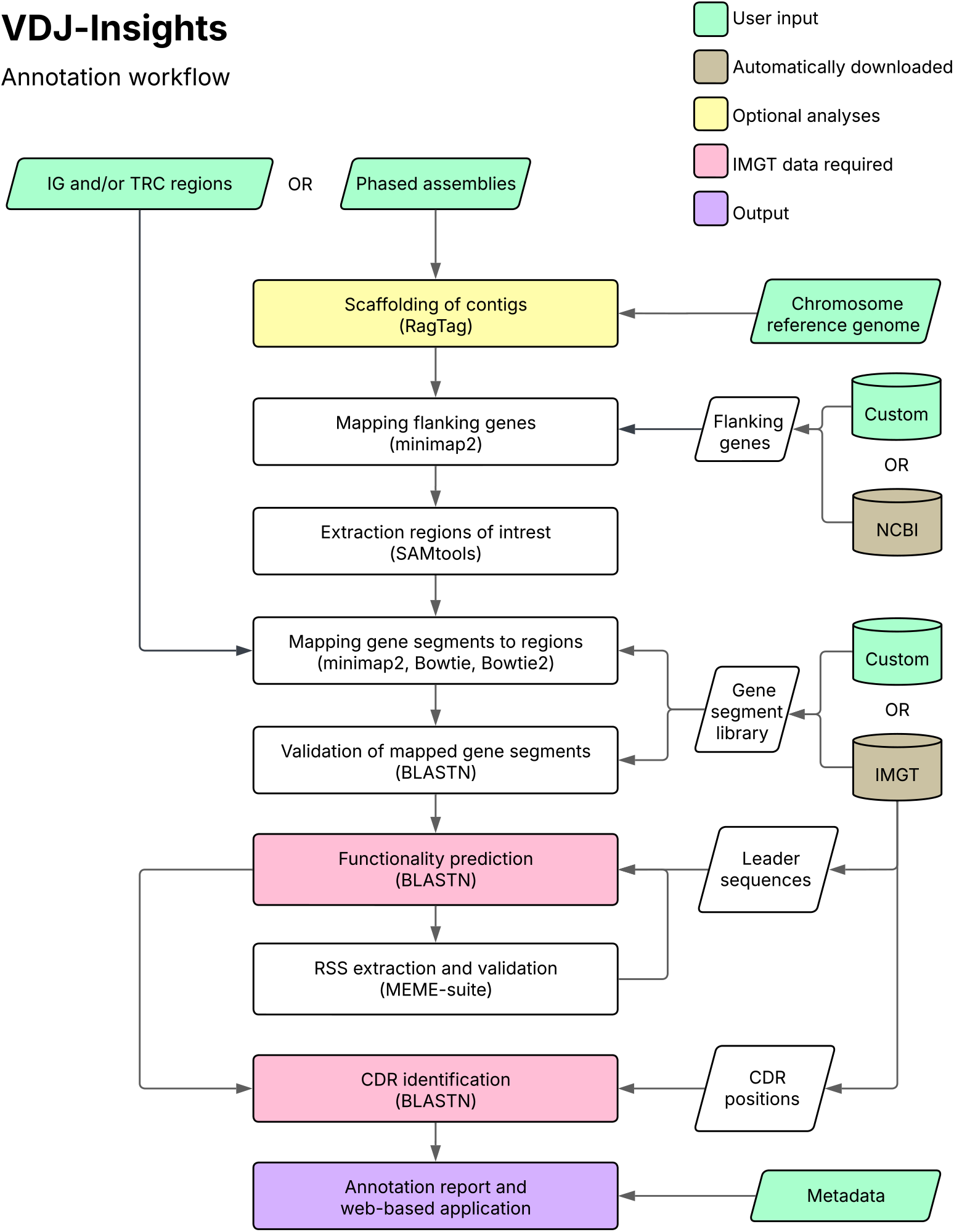
Workflow of the VDJ-Insights IG and TCR annotation. Different colours indicate user-provided input, input automatically retrieved by VDJ-Insights, optional analyses, and analyses requiring IMGT data. For each step, the primary tools used are listed in parentheses.

The output of VDJ-Insights includes a comprehensive overview of the identified known and novel gene segments, their predicted functionality, and an assessment of the RSS, CDR1 and CDR2 sequences, which are compiled into an interactive web-based annotation report. This report facilitates downstream analyses of the extracted IG and TCR regions, including the comparison of gene segments across multiple samples and the evaluation of newly identified sequences.

### Benchmarking VDJ-Insights using IMGT-curated IG and TCR regions from human, non-human primates, and mice

To validate the accuracy and robustness of VDJ-Insights, we applied the tool to annotate IMGT-curated IG and TCR regions from a human, four gorillas (*Gorilla gorilla gorilla*), a rhesus macaque, and four house mouse strains (Suppl. Table S2) (22, 27, 33, 34). For each species, the latest gene segment libraries, comprising functional, open-reading frame (ORF), and pseudogene segments, were sourced from the IMGT database (release 2025-04-30) (43). This setup was also applied to run Digger, a recently published IG and TCR annotation tool, allowing a direct comparison of performance (29). The curated human dataset includes the IGH and all TCR loci, comprising a total of 399 gene segments according to IMGT annotation. Using VDJ-Insights, we correctly identified all but two gene segments (Fig. 2A). Additionally, the tool reported 16 extra gene segments absent from the IMGT annotation, which may represent false positives or gene segments not yet annotated by IMGT. These extra gene segments were predominantly D segments, which are inherently challenging to map accurately due to their short length, along with two V segments that were identical to known reference sequences. In comparison, Digger identified 387 of the 399 IMGT-curated segments and reported 185 additional gene segments, which may represent false positives, including 79 V segments (Fig. 2A). VDJ-Insights was subsequently applied to curated IG loci from gorilla, rhesus macaque, and mouse (IGH and IGL), where it also achieved higher annotation accuracy than Digger in all three species (Figs. 2B, 2C, and 2D and Table 2).

**Table 2.** Number of segments identified by VDJ-Insights and Digger, and their concordance with IMGT curated annotations.

| Species | IMGT curated annotations (#) | Tool | Gene segments identified (#) | In concordance with IMGT (#) | In concordance with IMGT (%) |
| --- | --- | --- | --- | --- | --- |
| Human | 399 | Digger | 572 | 387 | 96.99 |
|  |  | VDJ-Insights | 413 | 397 | 99.50 |
| Western lowland gorilla | 1209 | Digger | 1748 | 1135 | 93.88 |
|  |  | VDJ-Insights | 1227 | 1209 | 100.00 |
| Rhesus macaque | 504 | Digger | 687 | 483 | 95.83 |
|  |  | VDJ-Insights | 509 | 504 | 100.00 |
| House mouse | 1135 | Digger | 2573 | 1085 | 95.59 |
|  |  | VDJ-Insights | 1226 | 1126 | 99.21 |
| Total | 3247 | Digger | 5580 | 3090 | 95.16 |
|  |  | VDJ-Insights | 3375 | 3236 | 99.66 |

**Figure 2.**
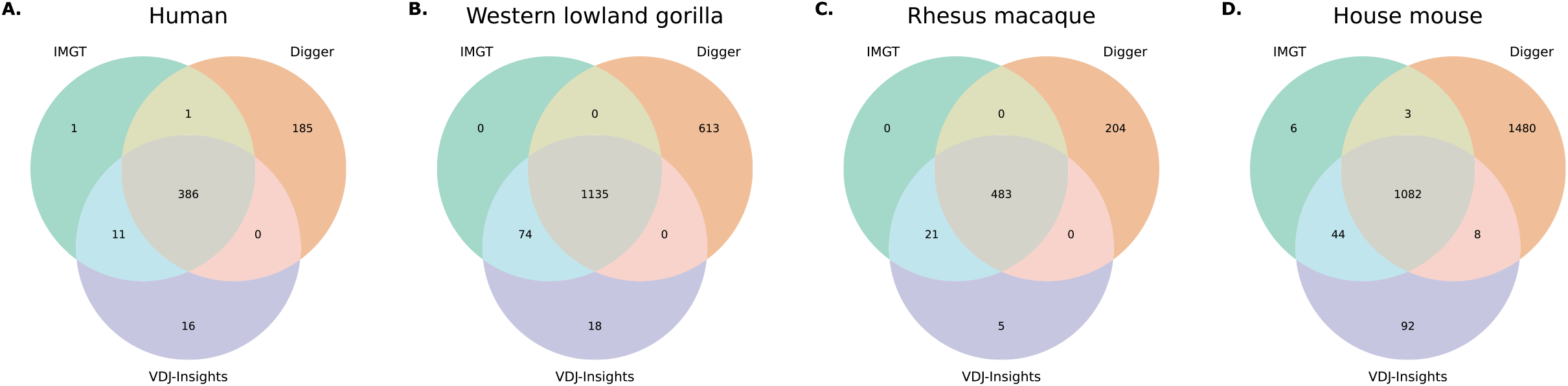
Comparison of gene segment annotations by VDJ-Insights, Digger, and IMGT. Venn diagrams depicting the overlap of annotated V, D, and J gene segments at both IG and TCR loci as identified by VDJ-Insights, Digger, and IMGT across the curated assemblies of a human (A), four gorillas (B), a rhesus macaque (C), and four mouse strains (D).

Next, we evaluated the accuracy of gene functionality classification as predicted by VDJ-Insights. The tool annotated 251 human gene segments as functional, 20 as ORFs, and 142 as pseudogenes, corresponding to concordance rates of 91.8%, 25,9%, and 98.4% with IMGT-curated classifications, respectively (Fig. 3 and Table 3). The observed discrepancies predominantly involved segments designated by IMGT as functional or ORFs (Suppl. Table S3), potentially reflecting the incorporation of additional information, such as experimental transcriptomic data, in the IMGT curation process (32). As compared to VDJ-Insights, Digger showed higher concordance with IMGT-curated functionality classifications for functional and ORF segments in humans, achieving agreement rates of 97.6% and 59.3%, respectively (Fig. 3 and Table 3). However, its accuracy in classifying pseudogene segments was lower, with a concordance rate of 59.8%. The higher concordance of Digger for functional and ORF classifications is likely a consequence of its default integration of IMGT-derived matrices, which incorporate curated information on gene functionality. A key limitation of this approach, however, is that Digger can only annotate in species for which such matrices are available. In contrast, the annotation process of VDJ-Insights does not depend on predefined references and is therefore applicable to a broader range of species (29). In the gorilla, rhesus macaque, and mouse datasets, functional and ORF classifications were largely consistent between the two tools, with Digger achieving higher concordance on functionality and ORF classification, whereas VDJ-Insights provided a more accurate annotation of pseudogene segments (Suppl. Fig. S1, Suppl. Table S4-6).

**Table 3.**
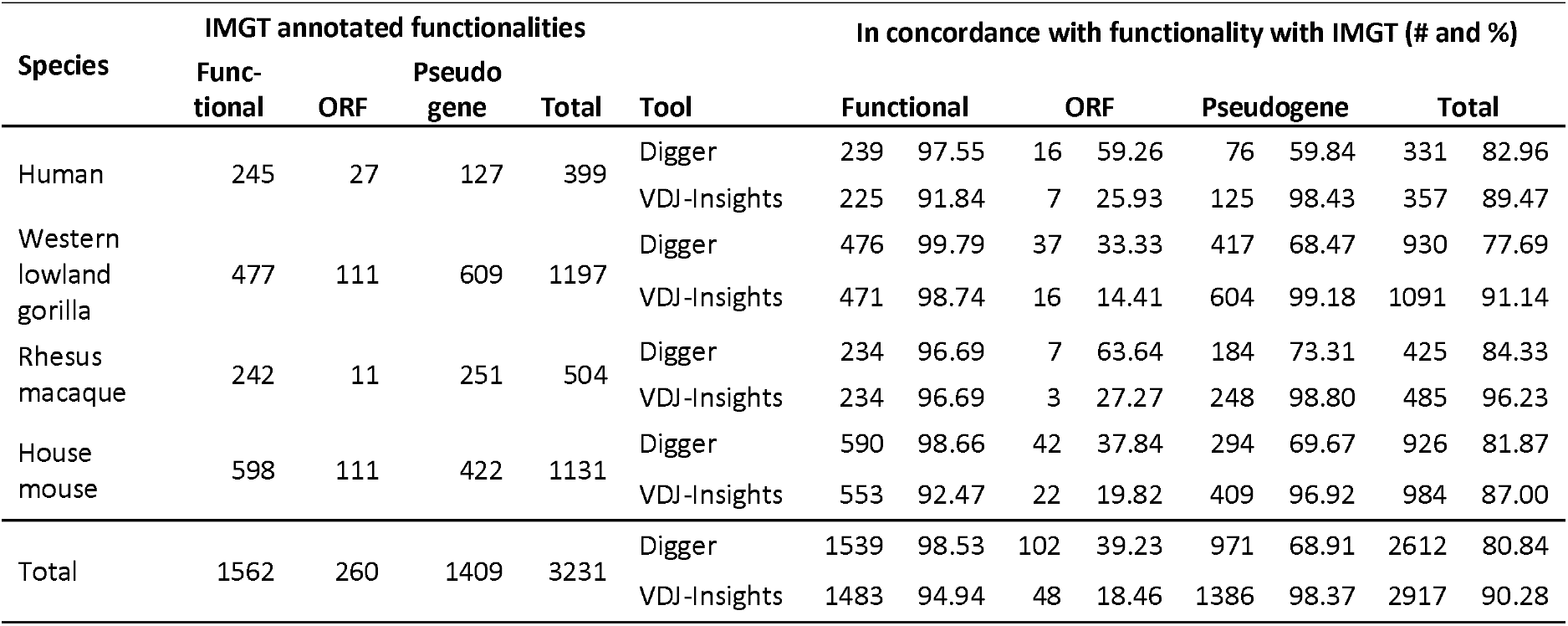
Functionality assessments of VDJ-Insights and Digger compared to IMGT curated annotations.

**Figure 3.**
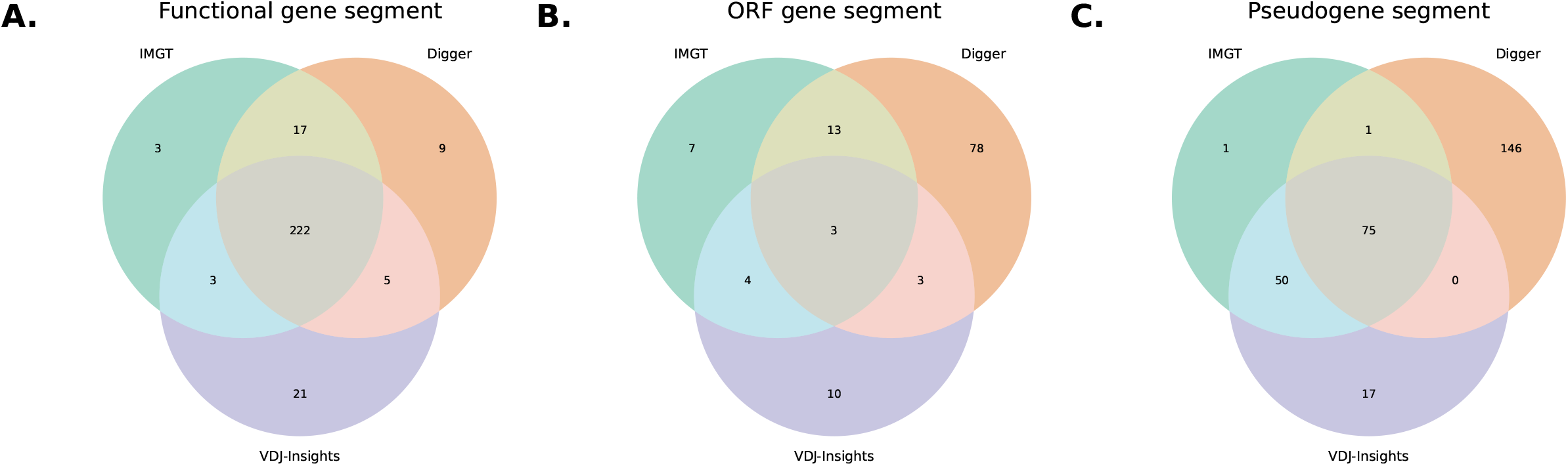
Comparison of functionality classification by VDJ-Insights, Digger, and IMGT. Venn diagrams illustrating the overlap in functionality classifications assigned by VDJ-Insights and Digger, compared to IMGT curated classifications, for human gene segments categorized as functional (A), open reading frame (ORF) (B), and pseudogenes (C).

**Figure 4.**
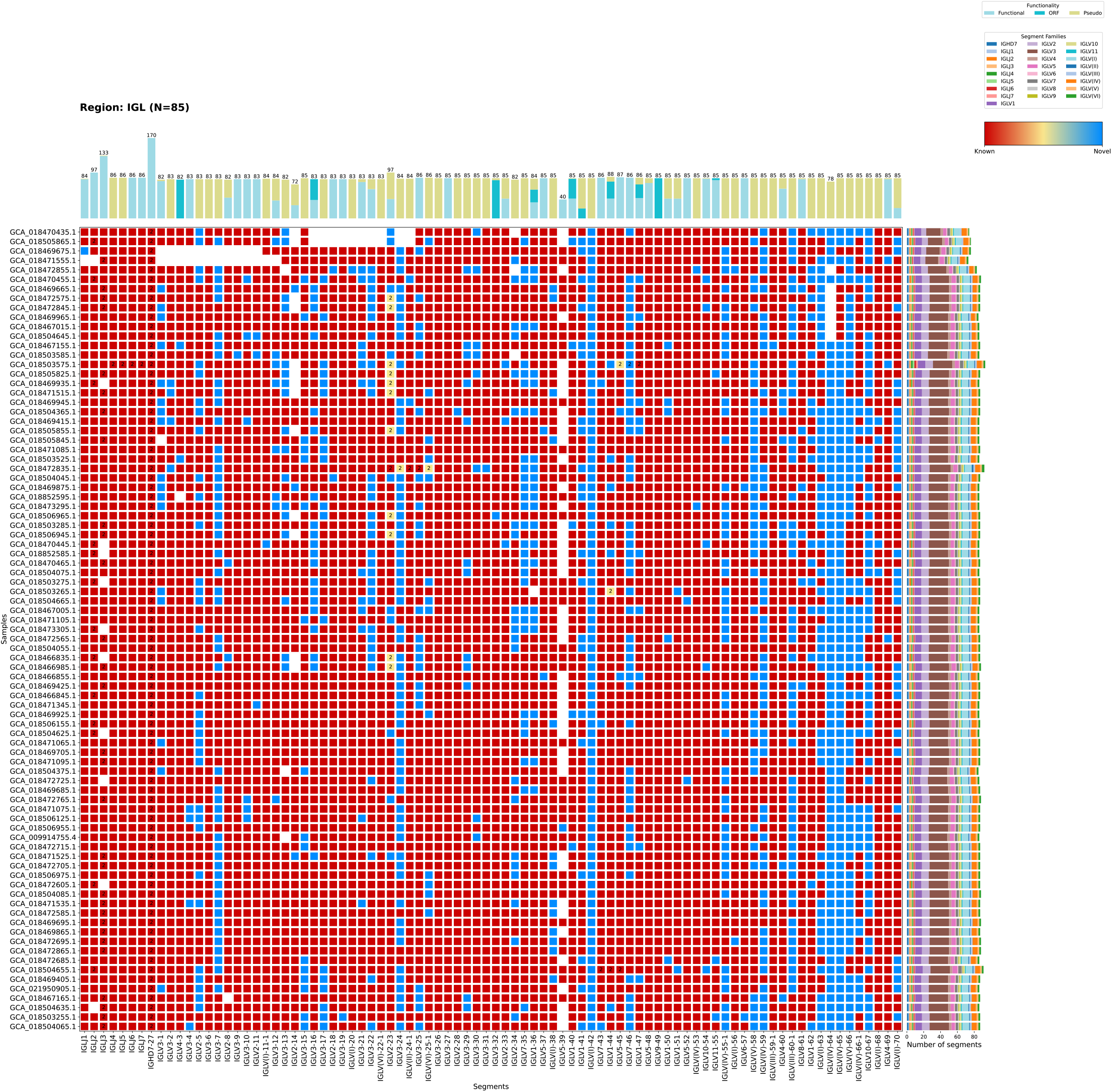
Visualization of gene segment presence across haplotypes in the IGL region. Each box indicates the presence (coloured) or absence (blank) of a specific gene segment per haplotype. Colours distinguish known alleles from novel ones based on existing gene segment libraries. Numbers within the boxes represent the number of duplicated segments. If multiple duplications are present, intermediate shading indicates the proportion of known versus novel alleles. The side chart illustrates the presence of gene segment families per haplotype, and the top chart summarizes the number of segments classified as functional, ORF or non-functional by VDJ-Insights.

Overall, we validated the accuracy and robustness of VDJ-Insights in identifying gene segments and classifying their functionality with high concordance to curated references. The reference-free approach positions VDJ-Insights as a valuable tool for extending IG and TCR region analyses beyond model-species with well-curated gene libraries.

### Characterisation of IG and TCR regions in the Human Pangenome dataset

The performance and computational efficiency of VDJ-Insights on large-scale datasets was evaluated by characterising the IG and TCR regions across 47 phased diploid human assemblies obtained from the Human Pan-Genome Reference Consortium (HPRC) and the Telomere-to-Telomere reference genome (T2T) of the CHM13 cell line (35, 44).

Although VDJ-Insights successfully identified the flanking genes in all samples, not all IG and TCR regions appeared to be completely or correctly assembled (Suppl. Fig. S2). This was most evident at the IGH and IGK loci, for which only three and four complete regions were identified, respectively. The difficulty in assembling these loci likely arises from their proximity to centromeric and telomeric regions of the chromosomes, which are known to pose challenges for genome assembly (35, 45, 46). Consequently, the relevant regions appear to be fragmented over multiple contigs. To resolve this issue, a scaffolding option was incorporated into VDJ-Insights, using the T2T genome as a reference (44). This approach enabled the recovery of an additional 26 IG loci, thereby increasing the overall coverage of IG and TCR haplotypes (Suppl. Fig. S2). VDJ-Insights flags scaffolded regions containing multiple contigs in its annotation report, as these assemblies contain unresolved gaps within the IG and TCR loci and may not fully capture their complete gene segment content. In total, 392 complete regions from the HPRC and T2T datasets were successfully annotated, including 115 IG (IGH, IGK, and IGL) and 277 TCR (TRA-TRD, TRB, and TRG) regions.

Two complementary public databases are available for human V, D, and J gene segments: IMGT, comprising curated germline sequences, and VDJbase, which contains genomic sequences from over 200 individuals (9, 43). Although VDJ-Insights is configured to access the most recent IMGT gene segment library by default, users can optionally utilize custom libraries. For the annotation of IG regions in the HPRC datasets, we employed three gene segment libraries: IMGT, VDJbase, and a custom combined library. The IMGT library contained 1051 unique IG gene segment sequences (release 2025-04-30), while the VDJbase library comprises 980 unique IG gene segments (assessed 2025-04-30), with 347 segments shared between these libraries. A combined library was generated by retaining one representative for each shared IG sequence and including all library-specific sequences, resulting in a total of 1,684 unique gene segments.

Using the IMGT library, a total of 26,865 gene segments were annotated across the IGH, IGK, and IGL loci from the 94 HPRC haplotypes and the T2T genome. In comparison, the VDJbase library annotated 13,779 gene segments. Utilizing the combined library, VDJ-Insights annotated a total of 27,514 gene segments (Suppl. Fig. 3 and Table 4). Among these annotated gene segments, 1,428 were unique, of which 652 were novel. Of these novel segments, 222 were confirmed in multiple individuals, providing strong evidence for their authenticity. Cross-comparison of the different libraries showed that 13,123 gene segments were consistently identified by all three reference datasets, whereas 13,742 and 656 gene segments were exclusively annotated by the IMGT and VDJbase libraries, respectively. These findings highlight the importance of comprehensive reference libraries for enhancing the accuracy of gene segment annotation.

**Table 4.**
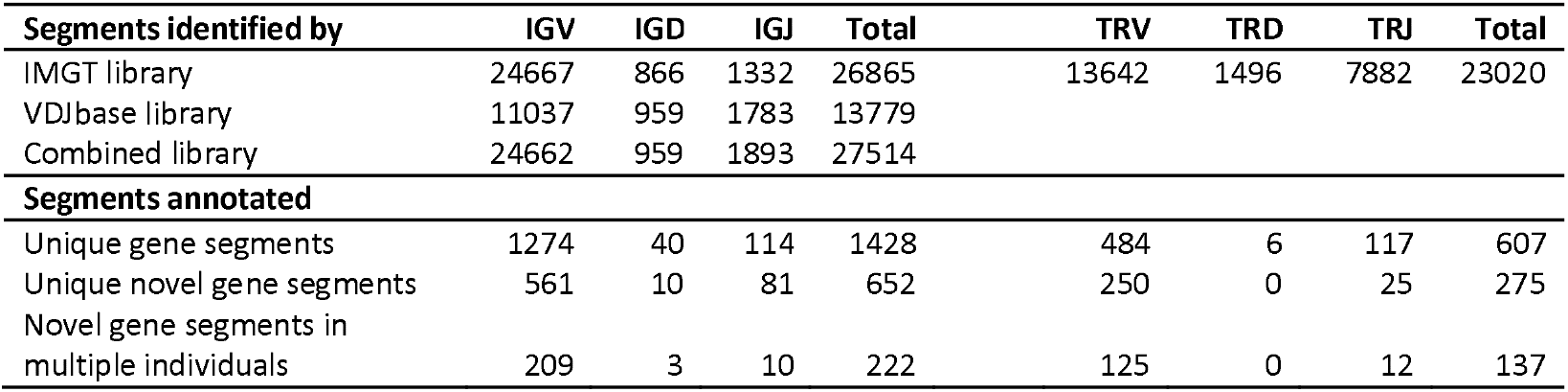
Number of gene segments annotated across the HPRC cohort for both the IG and TCR regions.

Since VDJbase exclusively includes V, D, and J gene segments for IG loci, TCR region annotation was performed solely using the IMGT library. In total, 23,020 TCR gene segments were annotated in the 95 human haplotypes, representing 607 unique alleles. Of these unique segments, 275 were identified as novel compared to the current IMGT reference. Furthermore, among these novel segments, 137 were documented in multiple individuals, providing additional support for their validity. At a haplotype level, we detected identical TCRB and TCRG haplotypes among HPRC samples, although at low frequencies (Suppl. Figs. 4D and 4E). These identical haplotypes were mainly identified in individuals of the same ethnicity (35), suggesting the presence of conserved TCR configurations within specific populations. All 46 studied individuals were heterozygous at their IG and TCR loci, except for one individual (HG01928, haplotypes GCA_018472695.1 and GCA_018472705.1), who was homozygous for the TCRG locus.

In addition to gene segment annotation, VDJ-Insights automatically identifies the RSS, as well as the CDR1 and CDR2 for the annotated IG and TCR gene segments. To assess nucleotide variation, sequence motifs were generated for each of these features across all IG and TCR loci. Analysis of the RSS of functional gene segments revealed highly conserved motifs across loci (Fig. 5), consistent with the conserved nature of the RAG1 and RAG2 recombinase proteins that mediate V(D)J recombination (47). An exception to these conserved motives was observed in the TRDD segments, likely due to annotation challenges associated with their short segment length. This may result in false positives that obscure true motif patterns. Other loci showed position-specific variability within otherwise conserved motifs. For example, the fourth position of the V-nonamer sequence displayed greater variability across all three IG loci, suggesting that this position may play a less critical role in RAG complex binding. As expected, the CDR regions exhibit substantial sequence variation (Fig. 6), consistent with their central role in forming the antigen-binding sites and binding of IG and TCR receptors to the major histocompatibility complex (MHC) (48). The CDR sequences were identified using IMGT-derived reference coordinates, resulting in the annotation of 757 sequences for IG loci and 390 sequences for TCR loci. Although not all gene segments in the IMGT database include annotated CDR regions, the majority of functional and ORF-classified V segments do contain this information. In contrast, limited CDR sequences are documented for the TRD and TRG loci, largely due to their relatively small number of V gene segments. Sequence motif analysis of the CDRs displayed extensive variability, with no universally conserved amino acid positions. Nevertheless, certain residues appeared with increased frequency at specific positions. For example, IGH-CDR1 frequently begins with glycine, while TRBV-CDR1 often includes a histidine. Additionally, the length of CDRs varied extensively, exemplified by IGHV-CDR2, which ranged from 3 to 10 amino acids in length. This high degree of diversity in both sequence composition and length highlights the evolutionary importance of CDR variability in supporting the broad antigen recognition capabilities of the adaptive immune system.

**Figure 5.**
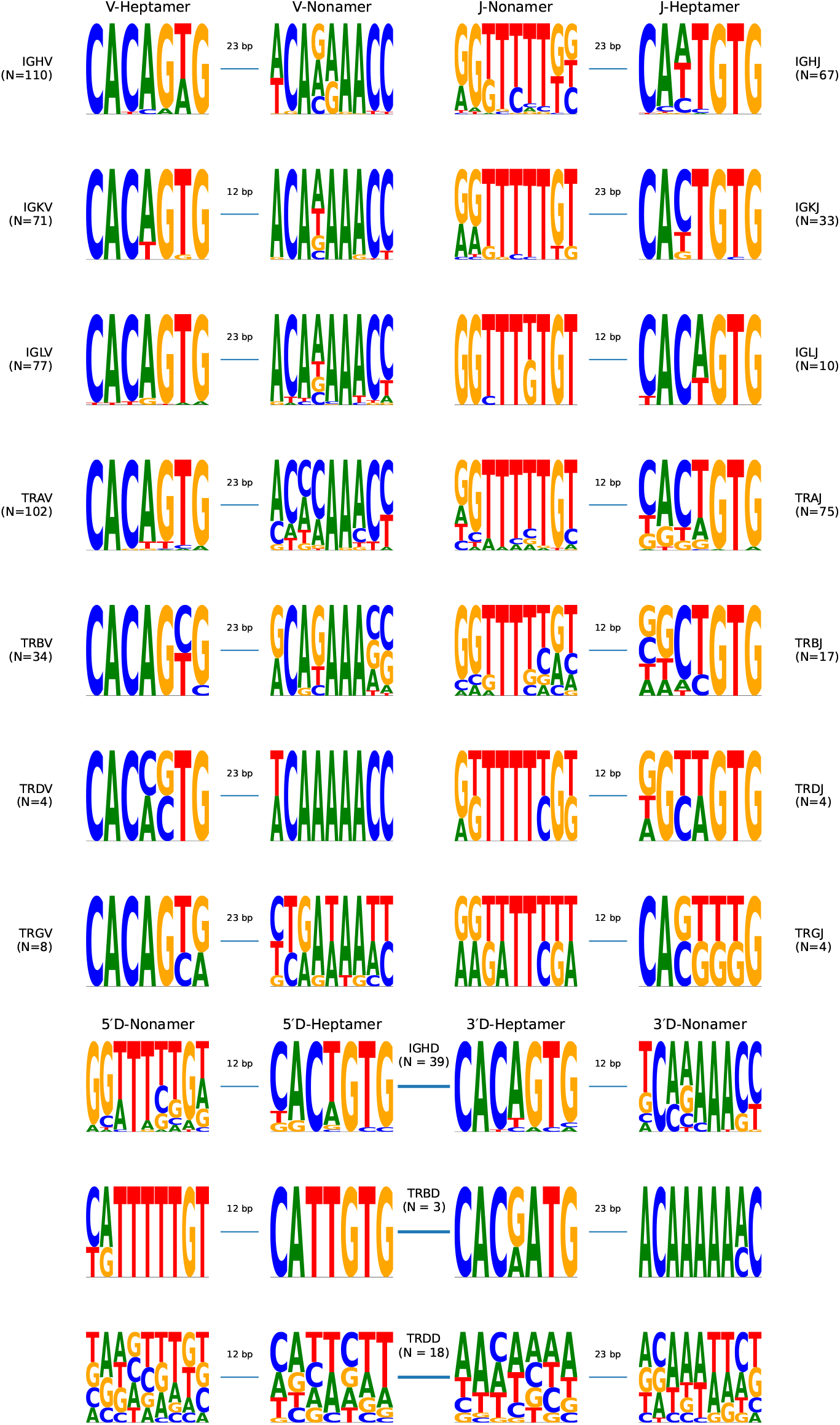
Recombination signal sequences (RSS) of V, D, and J gene segments identified across human IG and TCR loci. Each row represents one locus, and the height of each symbol indicates the relative frequency of the corresponding nucleotide at each position. Every gene segment is flanked by a heptamer, followed by a spacer (12 or 23 bps), and a nonamer. D segments display two RSS on both 5’ and 3’ sides. Only unique allele and RSS sequence combinations from functional gene segments (N) were included to generate these sequence motifs.

**Figure 6.**
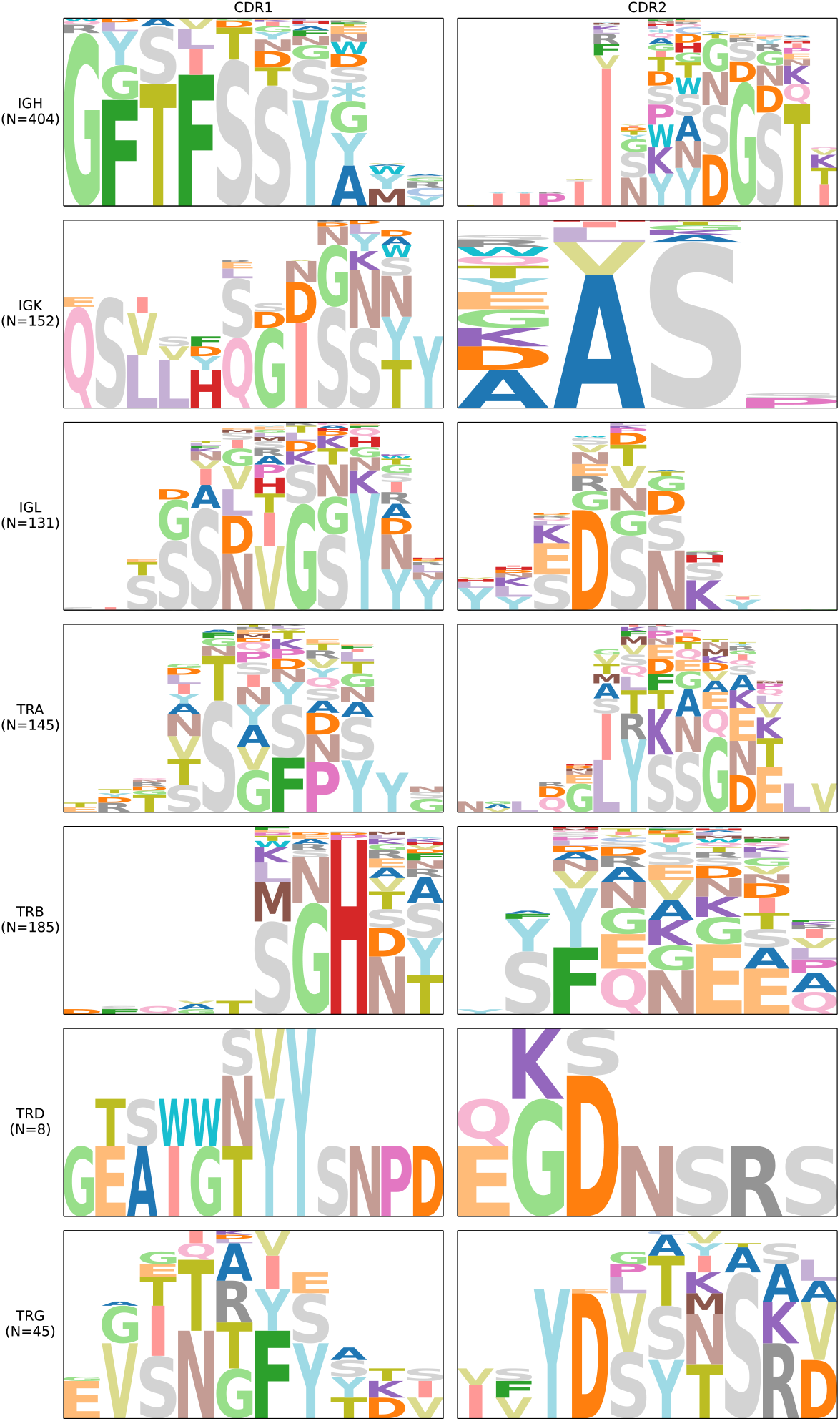
CDR1 and CDR2 sequence motifs across human IG and TCR loci. Each row represents a distinct locus, showing sequence logos for both CDR1 (left) and CDR2 (right). The height of each amino acid symbol indicates its relative frequency at that position within aligned sequences. Motifs were generated from alignments of CDR sequences of unique alleles (N) for each region. Positions where symbols do not reach the full height reflect alignment gaps, highlighting the variability in CDR lengths.

Overall, the in-depth characterization of IG and TCR immune regions in the HPRC cohort highlights the strength and robustness of our annotation tool. With the growing availability of large-scale genomic data, VDJ-Insights provides a comprehensive and benchmarked solution for automated IG and TCR annotation, advancing our understanding of the germline diversity that underlies adaptive immune responses.

## Discussion

Despite the central role of IG and TCR gene regions in the adaptive immune response, comprehensive germline characterization of these loci has historically remained limited due to their complexity (3). Recent advances in long-read and high-throughput sequencing technologies have substantially improved the potential to characterize these regions (4–8). However, the accurate identification and annotation of IG and TCR gene segments remains a challenging and time-intensive task, necessitating robust analytical tools and expert curation. VDJ-Insights was developed to address this challenge, providing a fast and reliable platform capable of annotating both known and novel gene segments across the highly polymorphic IG and TCR regions.

When benchmarked against expert-curated datasets from a human, gorillas, a rhesus macaque, and mice, VDJ-Insights successfully identified over 99% of the annotated segments (Table 2). VDJ-Insights also identified several additional gene segments absent from the curated references. Many of these were D gene segments, which are particularly difficult to annotate due to their short length, sometimes as small as eight base pairs. Among the few additional V segments, further investigation is warranted to determine whether these represent genuine novel alleles or potential false positives. VDJ-Insights also showed strong concordance in functionality classification with the reference annotations (Table 3). Discrepancies in functionality classification mostly involved segments labeled as ORFs. Several factors may account for these differences. For instance, while IMGT assigns a single functionality status to each allele, we can now demonstrate that variations in RSS or leader sequences can occur among identical allele sequences, potentially resulting in distinct functionality outcomes for the same gene segment. Moreover, IMGT’s integration of transcriptomic datasets, where available, further informs their functional assignments. As a result, functionality classification of annotation tools like VDJ-Insights, which rely exclusively on genomic sequence data and predefined rules, may occasionally diverge from IMGT’s more nuanced classifications.

During validation, we compared VDJ-Insights with Digger, a recently developed tool for IG and TCR gene annotation, which demonstrated slightly higher performance in predicting functional gene segments (29). This enhanced performance is likely due to Digger’s use of position weight matrices (PWMs) derived from IMGT-curated annotations. While this approach increases functional classification accuracy for well-characterized species, it limits the applicability of Digger to species or alleles represented in existing reference datasets. In contrast, VDJ-Insights is designed to minimize dependence on external reference databases for gene segment. This makes VDJ-Insights particularly well-suited for application to understudied or non-model organisms, including those lacking comprehensive IG or TCR gene annotations. Furthermore, VDJ-Insights is capable of processing multiple samples in parallel, enabling high-throughput analysis for large-scale genomic studies, such as pangenome assemblies and population-level cohorts. We demonstrated this scalability by annotating 94 haplotypes from the HPRC and the T2T human reference genome (35, 44), highlighting the efficiency and robustness of VDJ-Insights across extensive datasets. The accuracy of annotations, however, remains highly dependent on the quality of the underlying genome assemblies. This is especially true for loci such as IGH and IGK, which are located near telomeric and centromeric regions that are notoriously difficult to assemble. The extraction of the IGH locus is further complicated by the absence of a conserved flanking gene on one side, reducing the ability to define region boundaries. Even with high-quality HPRC assemblies, generated using a combination of Pacific Biosciences HiFi reads, Oxford Nanopore long reads, Bionano optical maps, and Hi-C Illumina short reads, several IG regions remained fragmented across multiple contigs (35). To address these challenges, VDJ-Insights provides the option to implement a reference-guided scaffolding strategy to improve contiguity of fragmented regions. This extra feature enhances the capacity to produce accurate and more complete annotations, even in fragmented assemblies.

In addition to the need for high-quality assemblies, our analysis emphasizes the critical role of a comprehensive and well-curated gene segment library in achieving accurate IG and TCR annotations.

This became evident during comparative evaluations using different reference libraries. Notably, when relying exclusively on the VDJbase library, we observed a highly incomplete annotation within the HPRC cohort (9). Utilizing larger and more inclusive gene segment libraries, not only improves detection sensitivity but also enhances the accuracy of functionality predictions. This is due to the adaptive modeling of RSS motifs using PWMs. As VDJ-Insights dynamically adjusts motif models based on the reference segments provided, richer libraries enable more robust functionality assessments. Moreover, VDJ-Insights accommodates post-analysis updates to reference libraries, allowing re-analysis with newly added or revised gene segments. This feature facilitates the annotation of previously unrecognized or misclassified segments, particularly as more novel alleles are discovered through large-scale sequencing efforts. However, it is essential to carefully evaluate the quality and reliability of added gene segments, especially for short segments such as D genes, which are inherently more prone to false-positive matches due to their limited sequence complexity. Our analysis of the IG and TCR loci across the HPRC cohort resulted in the annotation of 2,035 unique gene segments, of which 927 represent novel sequences not previously cataloged in IMGT or VDJbase. Among these novel segments, 359 were identified in multiple individuals, supporting their validity (Table 4). RSS motifs were extracted from all annotated segments and revealed a high degree of sequence conservation, consistent with their essential role in guiding V(D)J recombination. In parallel, CDR1 and CDR2 regions were annotated for 1,070 alleles using reference coordinates derived from IMGT. As expected, these regions displayed substantial sequence variability, which underlies their importance in directly recognizing antigens and engaging peptides presented by MHC molecules (48). Both RSS and CDR annotations offer valuable opportunities for downstream analyses. For example, variations in RSS motifs, in conjunction with segment positioning within the regions, have been linked to differences in V(D)J recombination frequency (49, 50). Similarly, availability of genomic CDR sequences enables investigations into somatic hypermutation and the functional evolution of the immune repertoire (48).

## Conclusions

VDJ⍰Insights provide a scalable and user⍰friendly software tool which accurately annotates IG and TCR regions from genomic assemblies. It identifies both known and novel gene segments, along with their associated RSS, CDR1, and CDR2 elements, predicts gene segments functionality, and facilitates downstream analyses with an interactive web-based interface. These features extend the utility of VDJ-Insights beyond gene segment annotation, offering a framework for functional immunogenetic as well as evolutionary comparative exploration at the individual, population, and species levels.

## Methods

### Details of the VDJ-Insights workflow

VDJ-Insights accepts one or more assembled whole genomes, phased contigs, or pre-extracted target regions from any species as input and performs a comprehensive annotation analysis of the IG and TCR loci. The pipeline is organized into modular components, each responsible for a specific task in the workflow. These modules are executed via a Python-based driver script, which calls the required software tools for each step. VDJ-Insights adheres to recommended thread and memory settings specified by the tools it integrates. It supports multiprocessing by distributing available resources across samples within the same module, while reserving a 10% memory buffer to prevent system overload and ensure stable performance. Optional modules can be activated using command-line flags, such as the scaffolding of phased contigs prior to the identification and extraction of IG and TCR loci.

1. *Scaffolding of phased contigs (optional)*. Discontinuous assemblies, often resulting from low sequencing coverage or extended homozygous regions, can cause the IG and TCR loci to be split across multiple contigs. To address this, VDJ-Insights offers an optional scaffolding step using RagTag (v2.1.0) (38) with default settings, which aims to reconstruct more complete IG and TCR regions by aligning contigs to a reference sequence. This step can be activated using the `-S` flag in the command line, followed by the path to a reference FASTA file for scaffolding. Any immune loci annotated from scaffolded sequences will be flagged in the final annotation report, as these may still lack complete gene segment representation. VDJ-Insights also generates scaffold visualizations, showing which contigs were joined and how many gene segments were identified on each contig within the scaffold.
2. *Identification and extraction of IG and TCR regions*. To ensure annotation of the complete IG and TCR loci, VDJ-Insights first locates these regions using conserved flanking genes. For several model species, such as human, rhesus macaque, and mouse, these flanking genes are preconfigured as default settings within the tool (Suppl. Table S1). For other species, users can specify custom flanking genes via the command line in JSON format (e.g., `-f ‘{“IGH”: [“PACS2”, “-”], “IGK”: [“RPIA”, “PAX8”], “IGL”: [“GANZ”, “TOP3B”]}’`). If a locus is located near a telomeric chromosome end, as is the case with the human IGH region on chromosome 14, the absence of a distal flanking gene can be indicated using a hyphen (“-”). The sequences of all specified flanking genes are automatically retrieved using the NCBI Datasets command-line interface (CLI) (v15.25.0) (51). The downloaded flanking gene sequences are aligned to the input assembly using minimap2 (v2.29) with asm5 settings (41). Based on these alignments, the corresponding genomic regions are extracted for downstream annotation using SAMtools (v1.22) (52).
3. *IMGT scraping of gene segment libraries, leader sequences, and CDR sequences*. VDJ-Insights includes a custom Python-based scraping tool developed to retrieve data from the IMGT database (https://www.imgt.org). By using the `-s` flag followed by a species name (e.g., `-s “Homo sapiens”`), the script automatically downloads gene segment libraries, leader sequences, including L-Part1 and L-Part2, and CDR1 and CDR2 sequences. If leader or CDR sequences are not available for a given species, VDJ-Insight automatically omits functionality classification or CDR analysis.
4. *Annotation of V, D, and J gene segments*. By default, V, D, and J gene segment reference libraries are retrieved from the IMGT database. Alternatively, users may provide custom gene segment libraries in FASTA format (`-l` flag). Extracted IG and TCR regions were annotated by aligning the segment libraries using three independent mapping tools run in parallel: minimap2 (v2.29) (41) with parameters `-a -m 70`, Bowtie (v1.3.1) (40) with parameters `-k 5 -a -M 5 --strata`, and Bowtie2 (v2.5.4) (39) with parameters `--end-to-end --very-sensitive --score-min L,0,-0.5`. The resulting alignment coordinates were converted to BED format using bedtools (v2.31.1) (53) and used to extract the corresponding nucleotide sequences from the input assembly. To confirm the identity of mapped gene segments, all extracted sequences were re-aligned using the megablast algorithm in BLASTN (v2.14.1) (42). For sequences shorter than 50 bp, additional parameters were applied to improve sensitivity, including `-penalty -3`, `-reward 1`, `-gapopen 5`, `-gapextend 2`, and `-word_size 7`. BLAST output was parsed to retain only alignments with 100% query coverage. Gene segments with 100% identity to reference sequences in the library were classified as known. Segments containing mismatches, insertions, or deletions were classified as novel and subjected to further mutation analysis using BTOP (BLAST traceback operations) parsing (54) to identify SNPs and indels at the nucleotide level.
5. *Determination of gene segment functionality*. The functionality of identified V, D, and J gene segments was predicted using a defined set of criteria (Table 1). Briefly, after retrieving leader sequences, comprising L-PART1 and L-PART2, from the IMGT database, genomic flanking regions adjacent to annotated V and J gene segments were subsequently extracted based on coordinate data, accounting for strand orientation. To align the L-part sequences to these genomic regions, the `blastn-short` task from BLASTN (42) was employed with parameters optimized for short and high-sensitivity alignments: `-word_size 7 -reward 1 -penalty -2 -gapopen 5 -gapextend 2 -best_hit_overhang 0.1 - best_hit_score_edge 0.1`. Mapped coordinates from the BLAST output enabled reconstruction of full-length pre-mRNA transcripts by concatenating L-PART1, L-PART2, and the V-intron. Canonical splice motifs, donor (GT) and acceptor (AG) sites, were identified at intron-exon boundaries to validate splicing integrity. The reconstructed nucleotide sequences were then translated into protein using the standard genetic code, facilitating the evaluation of open reading frames (ORFs), identification of functional start codons, premature stop codons, and other conserved protein features. Finally, recombination signal sequence (RSS) annotations were integrated from the downstream step, enabling comprehensive functional classification of each gene segment.
6. *RSS analysis*. For each V, D, and J segment, flanking nucleotide sequences containing the RSS were extracted from the corresponding genomic FASTA files based on their strand orientation and known coordinates. Two sets of RSS sequences were generated: one containing all RSS candidates, and a second subset containing only RSS sequences from known gene segments classified as functional by IMGT. These latter sequences were used to build representative motif models. Motif discovery was performed using the MEME Suite (v5.5.8) (55). The set of known functional RSS sequences were input to MEME with the following parameters: `-dna -mod zoops -nmotifs 1` and a fixed motif width equal to the expected RSS length. To detect motif occurrences across all extracted sequences, including novel gene segments, FIMO (Find Individual Motif Occurrences) (56) was run using the generated motif model and all RSS sequences as input, with a search threshold of `--thresh 0.0001` to maximize specificity and predict the final functionality. Segments lacking valid RSS motifs were annotated as pseudogene (Table 1).
7. *CDR analysis*. For each receptor type (IG or TCR) and segment class (e.g., IGHV, TRBV) genomic sequences were downloaded. The nucleotide sequences for CDR1 and CDR2 regions were extracted from each entry based on fixed positional offsets within the genomic sequences. Specifically, CDR1 was extracted from positions 78–113 and CDR2 from 165–194. These positions are based on established IMGT numbering conventions and were adjusted to remove gap characters (57). To map CDR regions onto the target sequences, BLASTN (42) was used in short-read alignment mode (`-task blastn-short`). For each target, the top-scoring alignment was selected using a ranking strategy prioritizing query start position, alignment coverage, and match length.
8. *Annotation report*. To identify, filter, and annotate immune gene segments from high-throughput sequence alignments, a comprehensive data processing application is added to VDJ-Insights. This Flask application integrates alignment parsing, variant detection, gene segment classification, and standardized output generation (including BED, GTF, and Excel reports). The process distinguishes between known and novel gene segments and captures associated metadata for downstream interpretation including segment-based principal component analysis, dendrogram construction, and Venn diagram generation.

### Computational performance assessment

The computational efficiency of VDJ-Insights was evaluated by annotating a subset of 20 randomly selected human HPRC haplotypes using varying numbers of processing threads, ranging from 4 to 20. On a workstation equipped with 258 GB of RAM, VDJ-Insights completed IG and TCR region annotations within 20 to 40 minutes (Suppl. Fig. S5). The tool operates with a minimum of 4 threads and achieves optimal performance when run with approximately 12 threads.

### Validation and HPRC datasets

The annotation accuracy of VDJ-Insights was validated using expertly curated immunogenetic reference sequences from the IMGT database (Suppl. Table S2). This benchmarking set included five human IG and TR loci from the GRCh38 human reference genome (34), three IG regions from four Western lowland gorilla (*Gorilla gorilla gorilla*) genome assemblies (27), IG regions from the rhesus macaque (*Macaca mulatta*) reference genome (22), and the IGH and IGL loci of four laboratory house mouse (*Mus musculus*) strains, including C57BL/6J, BALB/cJ, DBA/2J, and NOD/SCID (33). Gene segment annotations and predicted functionality generated by VDJ-Insights were systematically compared to these IMGT-curated references using a custom validation script, which is publicly available via GitHub.

Following validation, the tool was applied to analyse a comprehensive dataset of 95 haploid genome assemblies. This included the initial batch of genomes released by the Human Pangenome Reference Consortium (HPRC) (release 1) (35) as well as the most recent assembly from the Telomere-to-Telomere (T2T) Consortium (34). All assemblies were processed in a single execution, leveraging the parallel computing capabilities of VDJ-Insights. Sample metadata corresponding to the HPRC assemblies were retrieved from the HPRC Data Explorer (https://data.humanpangenome.org/assemblies).

### Annotating V(D)J gene segments using Digger

To further benchmark VDJ-Insights, all IMGT-curated sequences from human, Western lowland gorilla, rhesus macaque, and house mouse were also analysed using Digger (v0.7.5) (29). To support RSS and leader sequence analyses, position weight matrices (PWMs) were first generated for the gorilla and mouse datasets by parsing IMGT-derived annotations using the `parse_imgt_annotations` function. For each species and region, multiple annotation CSV files were merged, and PWMs were calculated using the `calc_motifs` function. Next, gapped V segment reference libraries were downloaded from the IMGT website using `extract_refs`. For the rhesus macaque, gaps in the reference sequences were corrected using the `fix_macaque_gaps` function to ensure compatibility with Digger’s input requirements. Digger was then executed on the same genomic regions as analyzed by VDJ-Insights, using identical gene segment libraries. These libraries were, however, first portioned into separate V, D, and J libraries for each species and region, as is required by Digger. All regions across all species were analyzed in both sense and anti-sense orientations direction (e.g. `digger` [r`egion_fasta] -—locus [locus]` `-—species [species] -—v_ref [V-library] -—d_ref [D-library] -—j_ref [J-library] -—v_ref_gap [gapped-V-library] -—sense [forwards/reverse] [output]`). Next, the curated IMGT annotations were compared with the outcomes of VDJ-Insights and Digger using a custom script (available on GitHub). Annotations with overlapping or identical start and stop coordinates were considered concordant across IMGT, VDJ-Insights, and Digger.

### RSS and CDR feature visualization

The RSS and CDR regions were analyzed using two custom scripts. For the RSS analyses, only unique allele and corresponding RSS combinations of functional gene segments were used. Sequence motifs were generated for each region using the Python package Logomaker (v0.8.4) (58).

For the CDR analyses, CDR1 and CDR2 regions were extracted from all unique alleles. The corresponding nucleotide sequences were translated to amino acids sequences, which were then aligned using Clustal-Omega (v1.2.4) (59). Subsequently, sequence motifs were generated from the aligned amino acid sequences using Logomaker (58).

## Declarations

### Ethics approval and consent to participate

Ethical approval is not applicable for this article

### Consent for publication

Not applicable.

### Availability of data and materials

The datasets generated during the current study are available in the VDJ-insights repository, https://github.com/BPRC-Bioinfo. Additional intermediate datasets used and analysed during the current study are available from the corresponding author on reasonable request.

## Acknowledgment

We would like to acknowledge the Human Pangenome Reference Consortium (BioProject ID: PRJNA730823) and its funder, the National Human Genome Research Institute (NHGRI).

## Authors’ contributions

J.B. conceived and designed the study. J.M. and S.J.M developed the VDJ-Insights software under the supervision of J.B., S.E.O., and G.N.L.. S.E.O. performed all data analyses, S.E.O. and S.J.M. were responsible for data visualization. S.E.O. and J.B. wrote the manuscript. N.G.G. contributed to methodological feedback. E.M.P. secured partial funding for the project. All authors reviewed, edited, and approved the final manuscript.

