## Supplementary material for "VDJ-Insights: simplifying the annotation of genomic IG and TCR regions": Suppl. Fig. S1

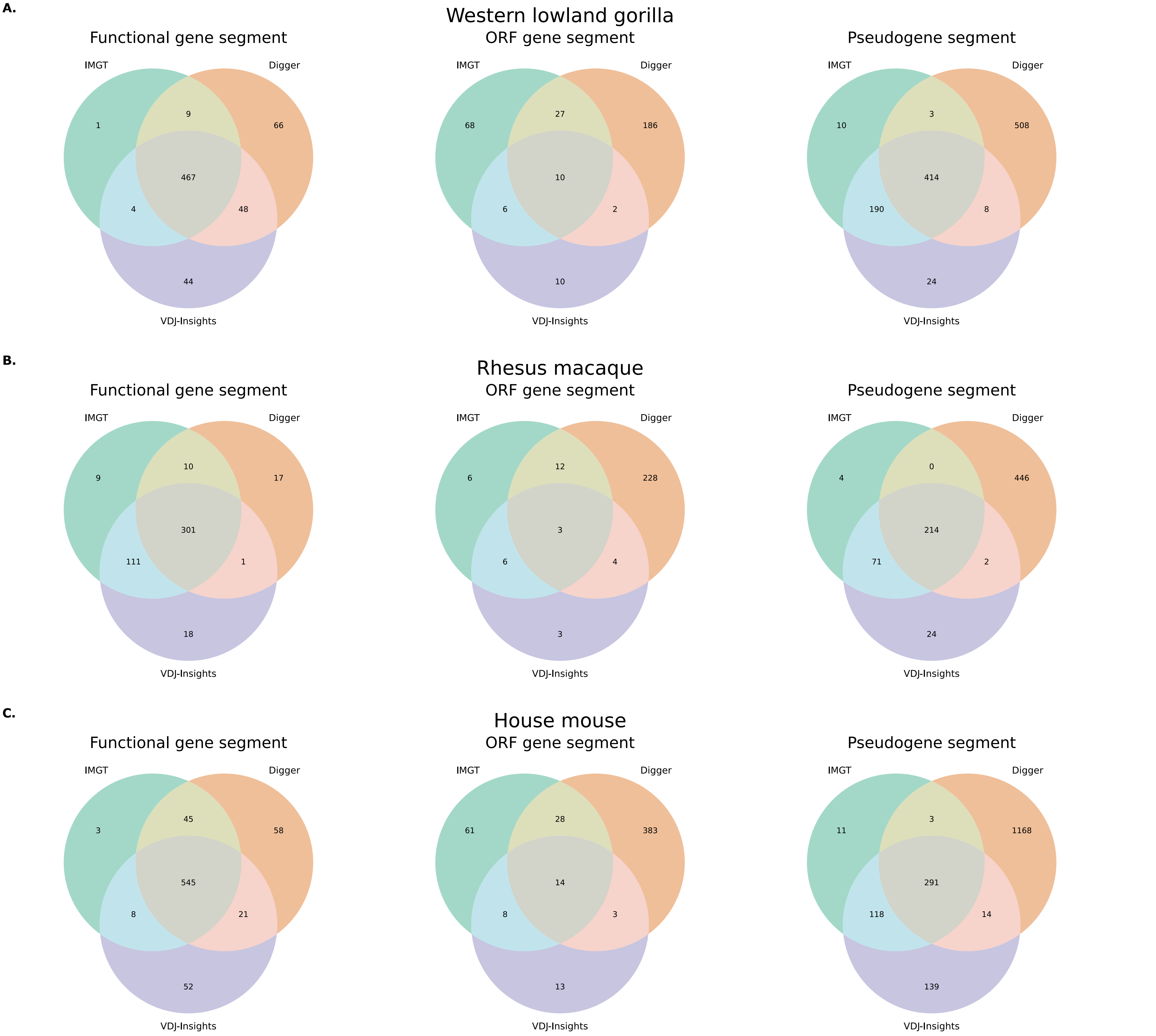


**Supplemental figure S1. Comparison of functionality classification by VDJ-Insights, Digger, and IMGT.** Venn diagrams illustrating the overlap in functionality classifications assigned by VDJ-Insights and Digger, compared to IMGT curated classifications, for gorillas (A), the rhesus macaque (B), and mice (C). Gene segments are categorized as functional, open reading frame (ORF), and pseudogenes.
