## Supplementary material for "VDJ-Insights: simplifying the annotation of genomic IG and TCR regions": Suppl. Fig. S2

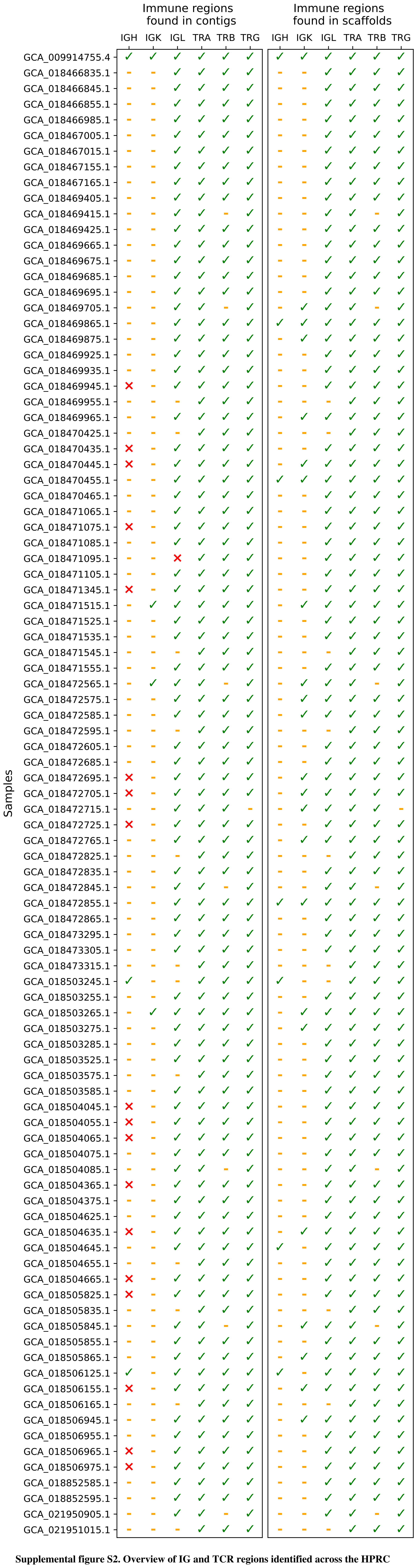


**Supplemental figure S2. Overview of IG and TCR regions identified across the HPRC cohort.** IG and TCR genomic regions were extracted and annotated from the whole-genome sequence contigs of the HPRC cohort (release 1, left) and the VDJ-Insights–scaffolded assemblies (right). The annotation status for each genomic region is depicted. A green check mark indicates successful extraction and annotation, defined as the locus being assembled onto a single contig or scaffold, with the number of annotated V, D, and J gene segments falling within the expected range as specified by IMGT (46, 61). An orange dash signifies an incomplete annotation, typically due to locus fragmentation across multiple contigs or annotation of fewer than 20% of the expected gene segments. A red cross denotes complete absence of annotated gene segments in the given region.
