## Supplementary material for "VDJ-Insights: simplifying the annotation of genomic IG and TCR regions": Suppl. Fig. S3

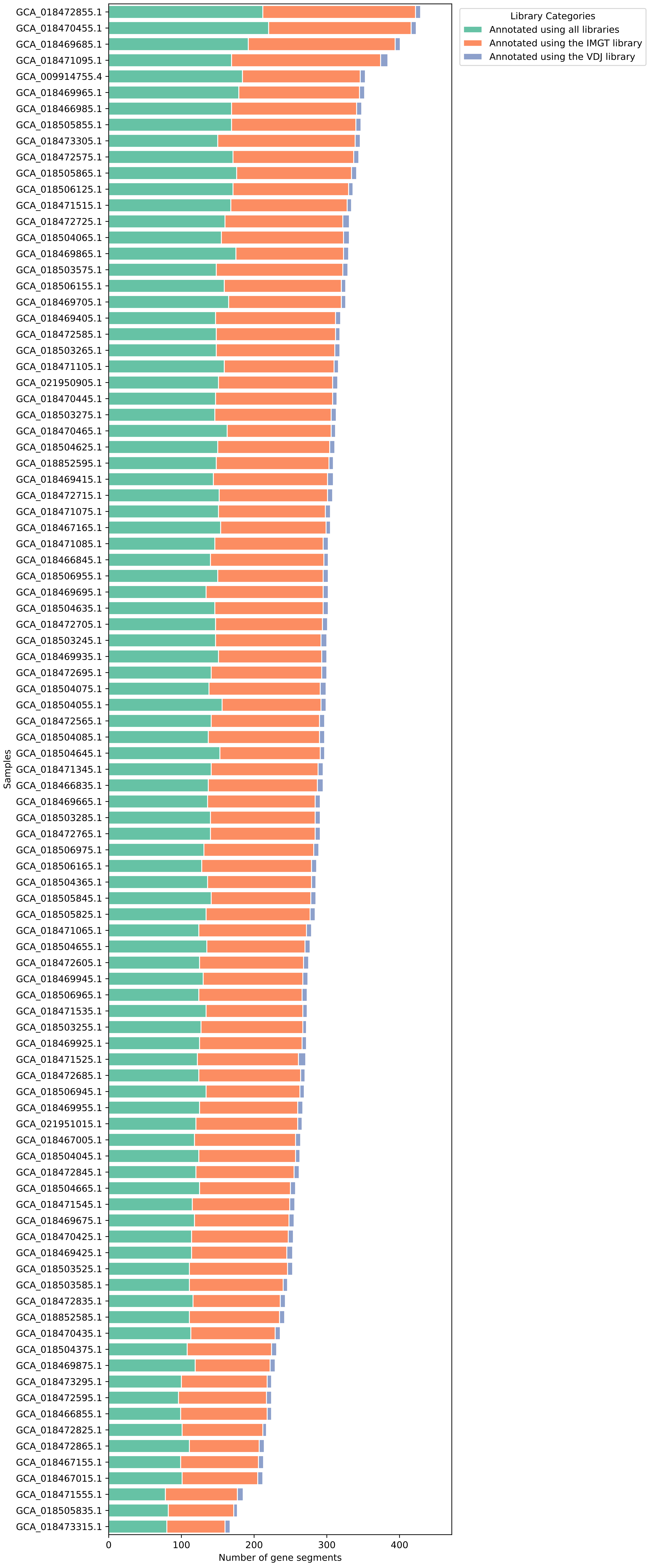


**Supplemental figure S3. Number of gene segments annotated using different reference libraries.** The bar chart illustrates how many gene segments were identified per haplotype, with colours indicating detection by the IMGT library, VDJbase library, or a combined library.
