## Supplementary material for "VDJ-Insights: simplifying the annotation of genomic IG and TCR regions": Suppl. Fig. S4

**Supplemental figure S4. Visualization of gene segment presence across haplotypes in different IG and TCR regions.** Each subplot (A-E) shows a different IG or TCR region. The boxes indicate the presence (coloured) or absence (blank) of a specific gene segment per haplotype. Colours distinguish known alleles from novel ones based on existing gene segment libraries. Numbers within the boxes represent the number of duplicated segments. If multiple duplications are present, intermediate shading indicates the proportion of known versus novel alleles. The side chart illustrates the presence of gene segment families per haplotype, and the top chart summarizes the number of segments classified as functional, ORF or non-functional by VDJ-Insights.

**A.**

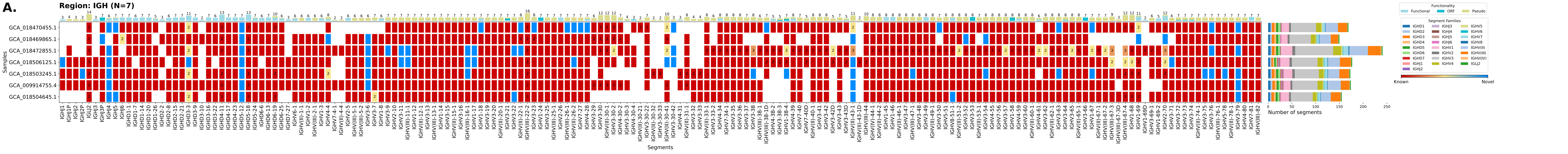

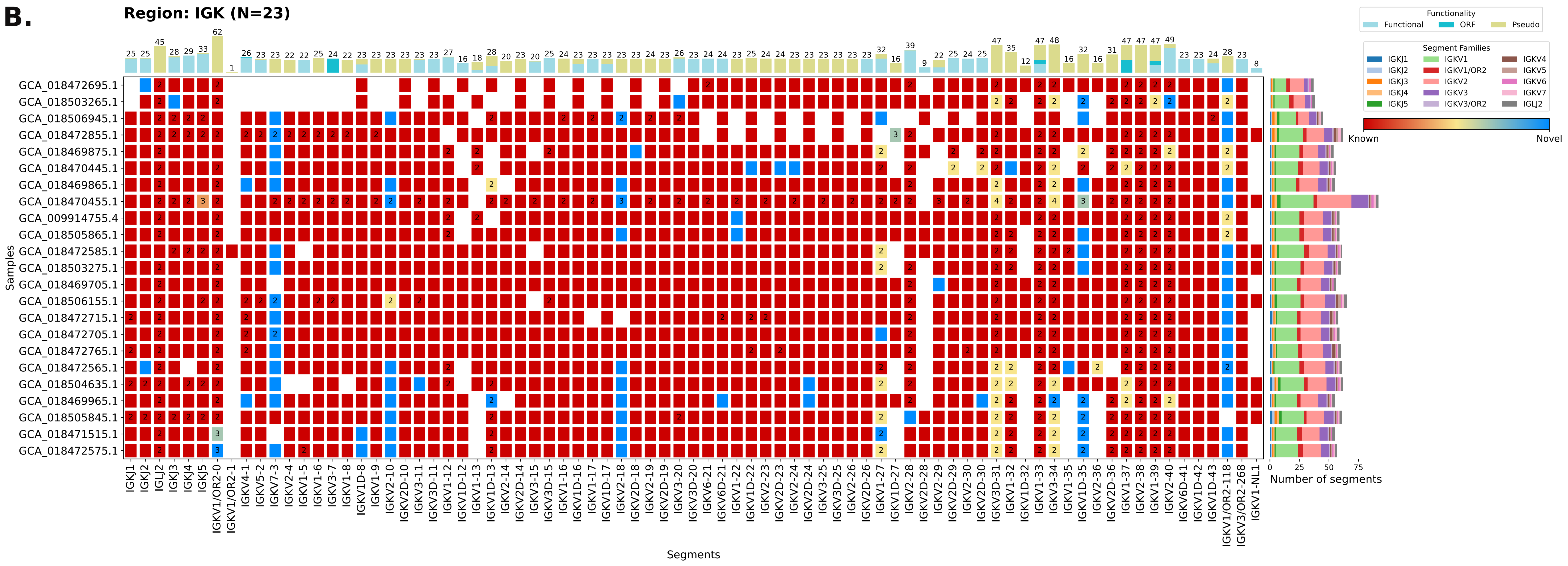
B.
Region: IGK (N=23)
Functionality
Functional
ORF
Pseudo
Segment Families
IGKJ1
IGKJ2
IGKJ3
IGKJ4
IGKJ5
IGKV1
IGKV1/OR2
IGKV2
IGKV3
IGKV3/OR2
IGKV4
IGKV5
IGKV6
IGKV7
IGLJ2
Known
Novel
Samples
GCA_018472695.1
GCA_018503265.1
GCA_018506945.1
GCA_018472855.1
GCA_018469875.1
GCA_018470445.1
GCA_018469865.1
GCA_018470455.1
GCA_009914755.4
GCA_018505865.1
GCA_018472585.1
GCA_018503275.1
GCA_018469705.1
GCA_018506155.1
GCA_018472715.1
GCA_018472705.1
GCA_018472765.1
GCA_018472565.1
GCA_018504635.1
GCA_018469965.1
GCA_018505845.1
GCA_018471515.1
GCA_018472575.1
IGKJ1
IGKJ2
IGLJ2
IGKJ3
IGKJ4
IGKJ5
IGKV1/OR2-0
IGKV1/OR2-1
IGKV4-1
IGKV5-2
IGKV7-3
IGKV2-4
IGKV1-5
IGKV1-6
IGKV3-7
IGKV1-8
IGKV1D-8
IGKV1-9
IGKV2-10
IGKV2D-10
IGKV3-11
IGKV3D-11
IGKV1-12
IGKV1D-12
IGKV1-13
IGKV1D-13
IGKV2-14
IGKV2D-14
IGKV3-15
IGKV3D-15
IGKV1-16
IGKV1D-16
IGKV1-17
IGKV1D-17
IGKV2-18
IGKV2D-18
IGKV2-19
IGKV2D-19
IGKV3-20
IGKV3D-20
IGKV6-21
IGKV6D-21
IGKV1-22
IGKV1D-22
IGKV2-23
IGKV2D-23
IGKV2-24
IGKV2D-24
IGKV3-25
IGKV3D-25
IGKV2-26
IGKV2D-26
IGKV1-27
IGKV1D-27
IGKV2-28
IGKV2D-28
IGKV2-29
IGKV2D-29
IGKV2-30
IGKV2D-30
IGKV3D-31
IGKV1-32
IGKV1D-32
IGKV1-33
IGKV3-34
IGKV1-35
IGKV1D-35
IGKV2-36
IGKV2D-36
IGKV1-37
IGKV2-38
IGKV1-39
IGKV2-40
IGKV6D-41
IGKV1D-42
IGKV1D-43
IGKV1/OR2-118
IGKV3/OR2-268
IGKV1-NL1
Segments
0
25
50
75
Number of segments

C.

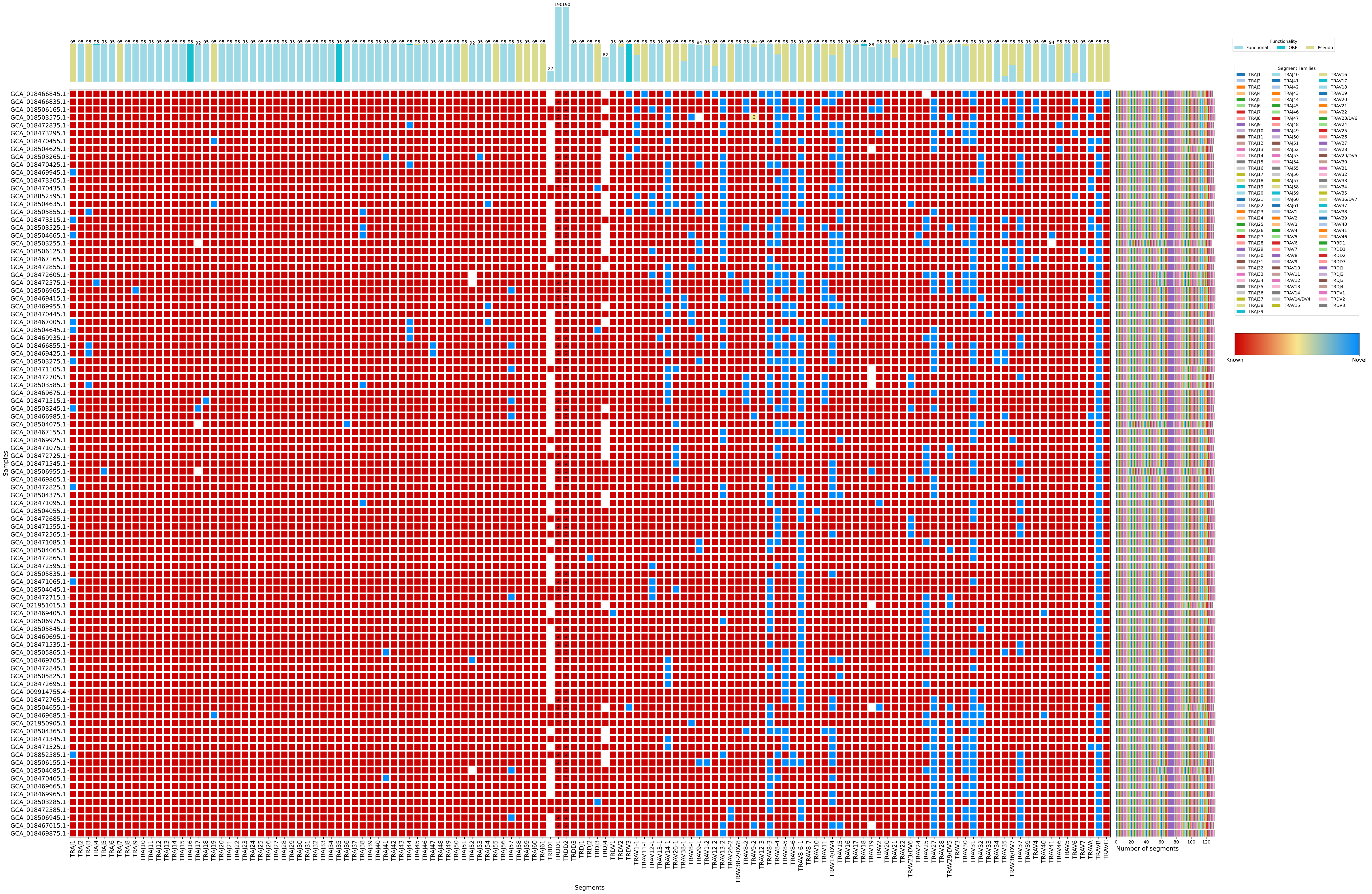

Region: TRA (N=95)
Samples
Segments
Number of segments
Functionality
Functional
ORF
Pseudo
Segment Families
Known
Novel

D.

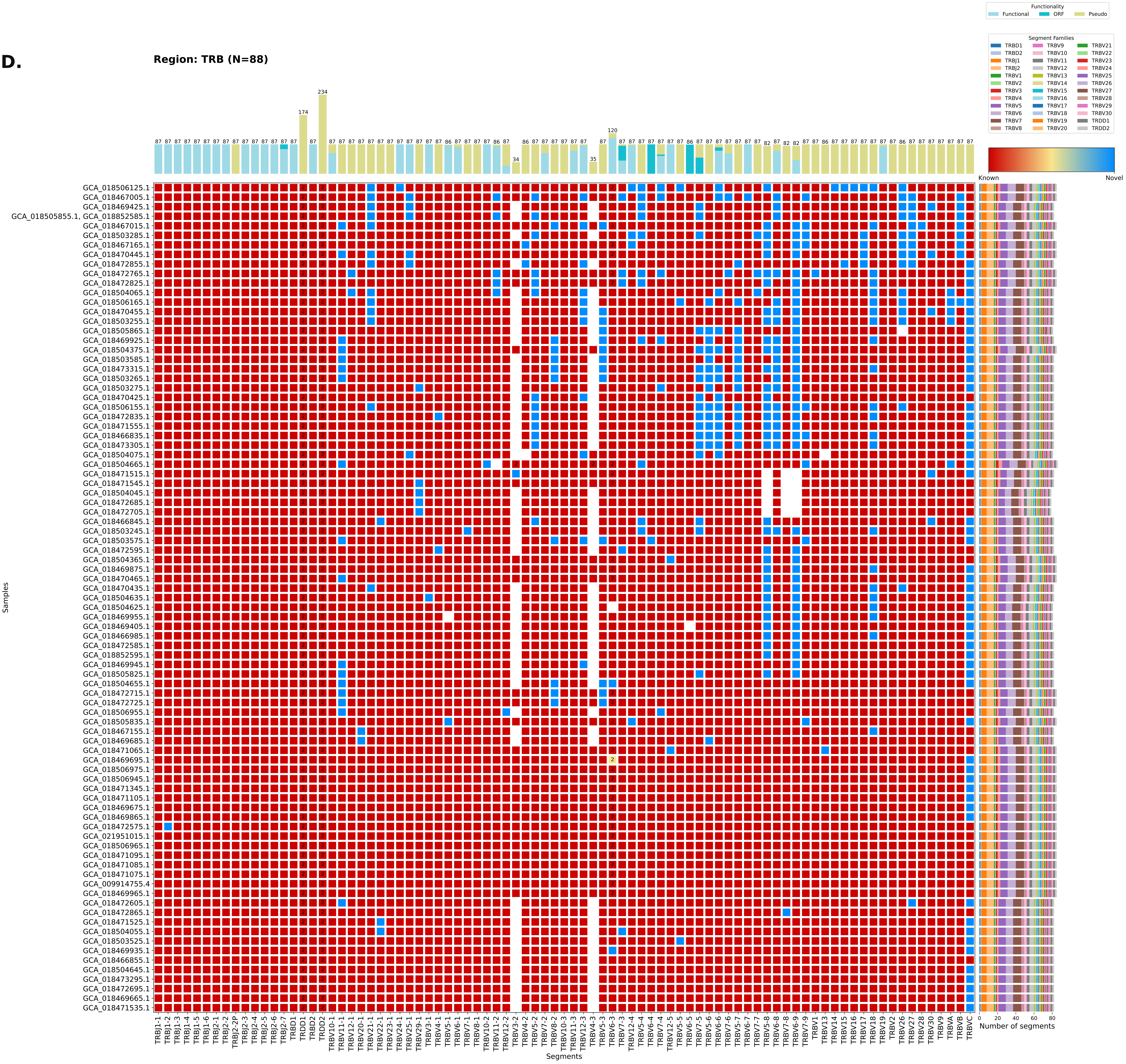

Region: TRB (N=88)
Functionality
Functional
ORF
Pseudo
Segment Families
Known
Novel
Samples
Segments
Number of segments

**E.**

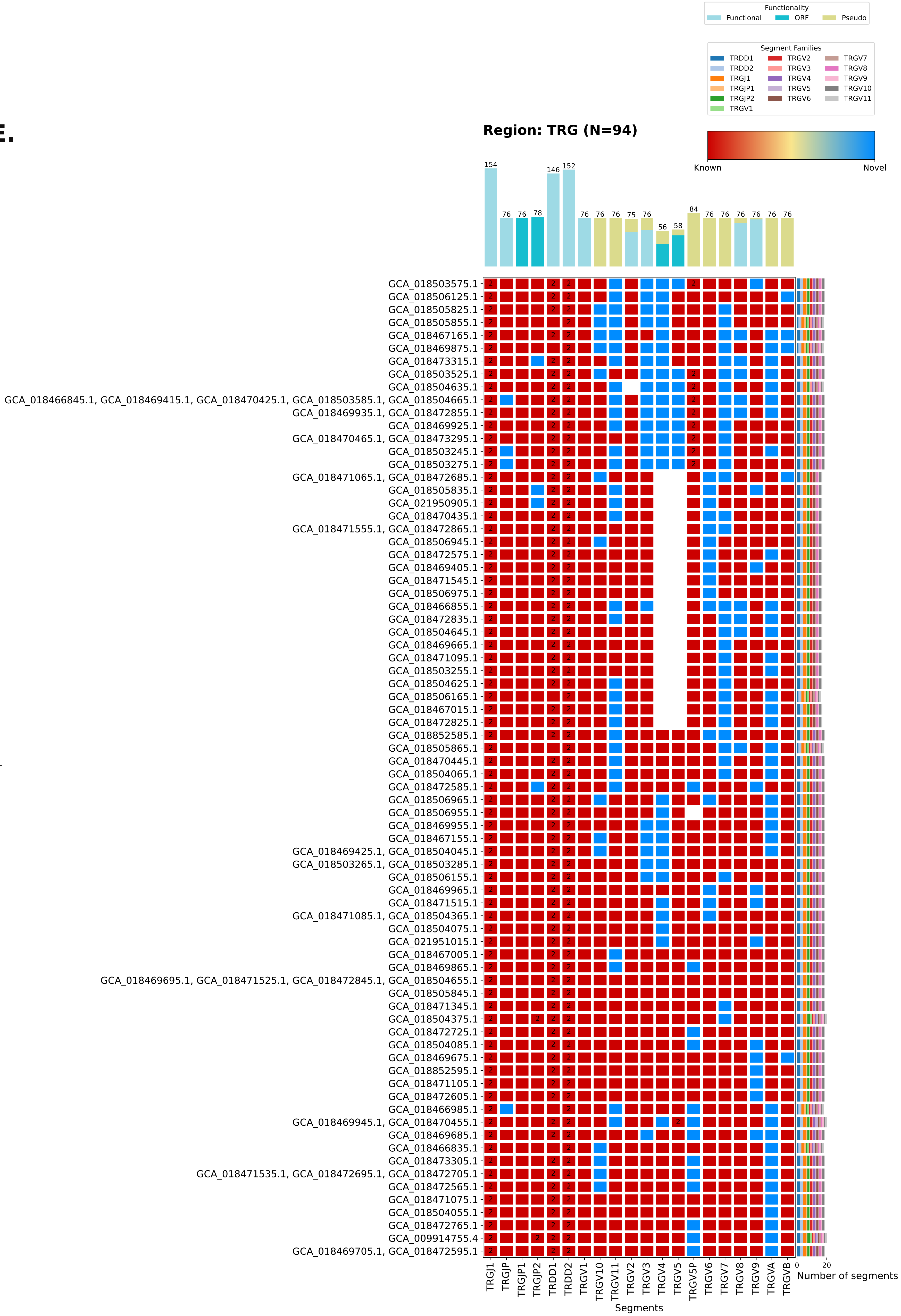
Functionality
Functional
ORF
Pseudo
Segment Families
TRDD1
TRDD2
TRGJ1
TRGJP1
TRGJP2
TRGV1
TRGV2
TRGV3
TRGV4
TRGV5
TRGV6
TRGV7
TRGV8
TRGV9
TRGV10
TRGV11
Region: TRG (N=94)
Known
Novel
154
76
76
78
146
152
76
76
76
75
76
56
58
84
76
76
76
76
76
76
GCA_018503575.1
GCA_018506125.1
GCA_018505825.1
GCA_018505855.1
GCA_018467165.1
GCA_018469875.1
GCA_018473315.1
GCA_018503525.1
GCA_018504635.1
GCA_018466845.1, GCA_018469415.1, GCA_018470425.1, GCA_018503585.1, GCA_018504665.1
GCA_018469935.1, GCA_018472855.1
GCA_018469925.1
GCA_018470465.1, GCA_018473295.1
GCA_018503245.1
GCA_018503275.1
GCA_018471065.1, GCA_018472685.1
GCA_018505835.1
GCA_021950905.1
GCA_018470435.1
GCA_018471555.1, GCA_018472865.1
GCA_018506945.1
GCA_018472575.1
GCA_018469405.1
GCA_018471545.1
GCA_018506975.1
GCA_018466855.1
GCA_018472835.1
GCA_018504645.1
GCA_018469665.1
GCA_018471095.1
GCA_018503255.1
GCA_018504625.1
GCA_018506165.1
GCA_018467015.1
GCA_018472825.1
GCA_018852585.1
GCA_018505865.1
GCA_018470445.1
GCA_018504065.1
GCA_018472585.1
GCA_018506965.1
GCA_018506955.1
GCA_018469955.1
GCA_018467155.1
GCA_018469425.1, GCA_018504045.1
GCA_018503265.1, GCA_018503285.1
GCA_018506155.1
GCA_018469965.1
GCA_018471515.1
GCA_018471085.1, GCA_018504365.1
GCA_018504075.1
GCA_021951015.1
GCA_018467005.1
GCA_018469865.1
GCA_018469695.1, GCA_018471525.1, GCA_018472845.1, GCA_018504655.1
GCA_018505845.1
GCA_018471345.1
GCA_018504375.1
GCA_018472725.1
GCA_018504085.1
GCA_018469675.1
GCA_018852595.1
GCA_018471105.1
GCA_018472605.1
GCA_018466985.1
GCA_018469945.1, GCA_018470455.1
GCA_018469685.1
GCA_018466835.1
GCA_018473305.1
GCA_018471535.1, GCA_018472695.1, GCA_018472705.1
GCA_018472565.1
GCA_018471075.1
GCA_018504055.1
GCA_018472765.1
GCA_009914755.4
GCA_018469705.1, GCA_018472595.1
Samples
TRGJ1
TRGJP
TRGJP1
TRGJP2
TRDD1
TRDD2
TRGV1
TRGV10
TRGV11
TRGV2
TRGV3
TRGV4
TRGV5
TRGV5P
TRGV6
TRGV7
TRGV8
TRGV9
TRGVA
TRGVB
Segments
0
20
Number of segments
