## Supplementary material for "VDJ-Insights: simplifying the annotation of genomic IG and TCR regions": Suppl. Fig. S5

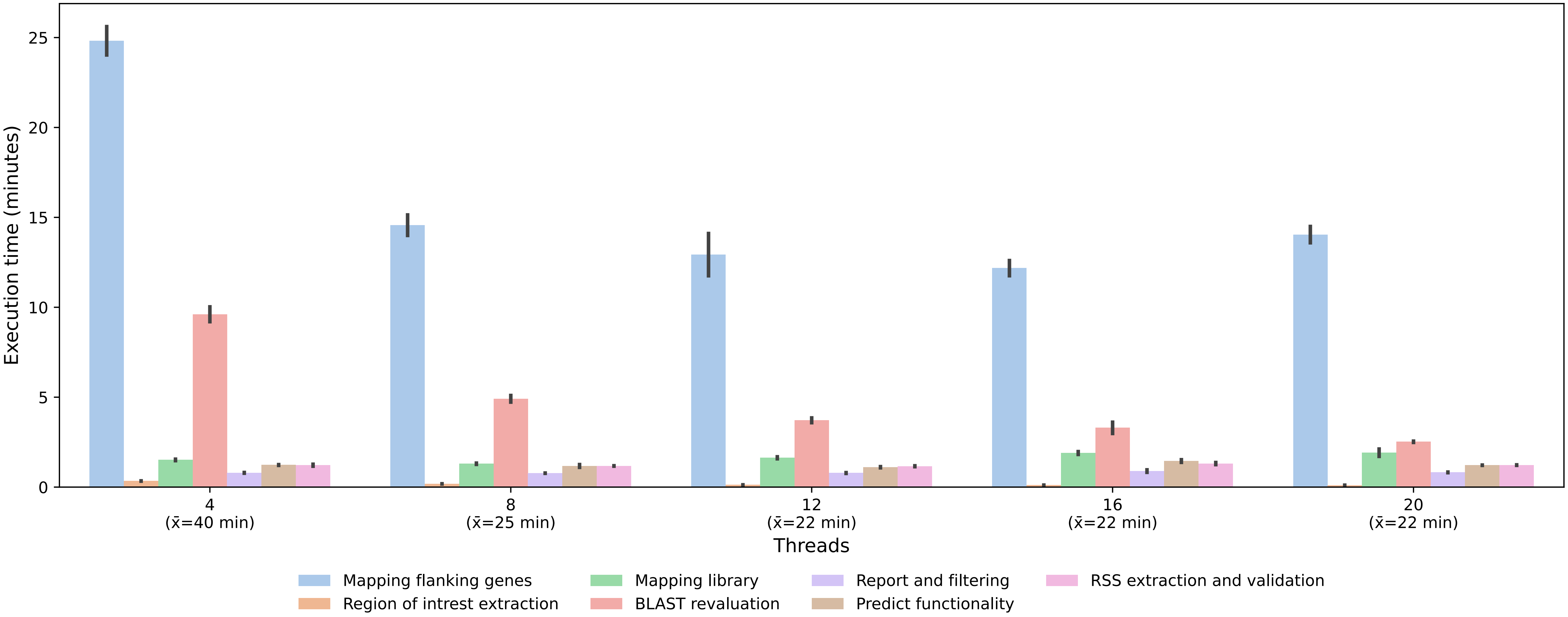


**Supplemental figure S5. Execution times of individual pipeline steps in VDJ-Insights for computational benchmarking.** The TCR regions from 20 randomly selected samples from the Human Pan-Genome Reference Consortium (HPRC) were extracted and annotated using their whole-genome assemblies. The analysis was performed on an AMD EPYC 7F72 24-Core Processor (utilizing 12 cores, 24 threads) with 258 GB RAM. Each colour corresponds to a distinct step in the VDJ-Insights pipeline, as detailed in the pipeline flowchart (Fig. 1). The x-axis indicates the number of threads utilized (4, 8, 12, 16, and 20). Results shown are averages ($\bar{x}$) across three independent runs, with error bars representing the standard deviation (SD).
